# Degradation of amyloid beta species by multi-copper oxidases

**DOI:** 10.1101/2023.07.02.547398

**Authors:** Jing Yang, Kathleen Ran, Wenzhe Ma, Lucy Chen, Cindy Chen, Can Zhang, Hui Ye, Ying Lu, Chongzhao Ran

**Affiliations:** Athinoula A. Martinos Center for Biomedical Imaging, Department of Radiology, Massachusetts General Hospital/Harvard Medical School, Room 2301, Building 149, Charlestown, Boston, Massachusetts 02129; School of Engineering, China Pharmaceutical University, Nanjing, 210009, China; Department of Systems Biology, Harvard Medical School, Boston, MA, United States; Genetics and Aging Research Unit, McCance Center for Brain Health, MassGeneral Institute for Neurodegenerative Disease, Department of Neurology, Massachusetts General Hospital and Harvard Medical School, Charlestown, MA, USA; Department of Biology, Loyola University Chicago, Chicago, IL, 60660

**Author notes:** Correspondence should be addressed to C.R. or Y.L.

## Abstract

Reduction of the production of amyloid beta (Aβ) species has been intensively investigated as potential therapeutic approaches for Alzheimer’s disease (AD). However, the degradation of Aβ species, another potential beneficial approach, has been far less explored. In this study, we discovered that ceruloplasmin (CP), an important multi-copper oxidase (MCO) in human blood, could degrade Aβ peptides. We also found that the presence of Vitamin C could enhance the degrading effect in a concentration-dependent manner. We then validated the CP-Aβ interaction using total internal reflection fluorescence (TIRF) microscopy, fluorescence photometer, and fluorescence polarization measurement. Based on the above discovery, we hypothesized that other MCOs had similar Aβ-degrading functions. Indeed, we found that other MCOs could induce Aβ degradation as well. Remarkably, we revealed that ascorbate oxidase (AO) had the strongest degrading effect among the tested MCOs. Using induced pluripotent stem (iPS) neuron cells, we observed that AO could rescue neuron toxicity which induced by Aβ oligomers. In addition, our electrophysiological analysis with brain slices suggested that AO could prevent an Aβ-induced deficit in synaptic transmission in the hippocampus. To the best of our knowledge, our report is the first to demonstrate that MCOs have a degrading function for peptides/proteins. Further investigations are warranted to explore the possible benefits of MCOs for future AD treatment.

## Introduction

The approval of antibody drugs (aducanumab and lecanemab) in the last two years demonstrates the potential of the amyloid cascade hypothesis to treat AD^1, 2^. Currently, most drugs that are based on the amyloid cascade hypothesis have been centered on reducing the production and accumulation of Aβ and preventing the aggregation of Aβs in brains ^3^. However, drugs that are capable of promoting the degradation of Aβs in brains and in peripheral systems have been paid much less attention ^4-9^. Notably, seeking protein degraders is one very active research area in drug discovery, and several small molecules based on proteolysis targeting chimera (PROTAC) have been investigated in clinical trials ^10 11 12^. We believe that seeking degraders for neurotoxic Aβs has great potential for AD drug discovery.

Multi-copper oxidases (MCO) are a diverse group of enzymes that catalyze the oxidation of various substrates, including polyphenols, aromatic polyamines, L-ascorbate, and metal ions such as iron (II) ^13-16^. Evidence suggested that several MCOs played an important role in lignocellulose degradation ^17, 18^. However, it has rarely been explored whether MCO can induce the degradation of peptides and proteins. In MCOs, copper plays an essential role in substrate binding and oxidation. All MCOs have four common copper ions in three distinct sites (Type I, II, and III copper centers), and all of the copper ions involve in coordination with at least two histidines ^19-21^. In our previous study, we demonstrated that the copper coordination with histidines (H6, H13/H14) of amyloid beta (Aβ) peptides could lead to the degradation of the Aβ peptides ^22^. In this regard, we speculated that MCOs could bind and degrade Aβ species as well.

In this report, indeed, we discovered that endogenous and exogenous multi-copper oxidases (MCO) could bind and degrade Aβ species in vitro solutions, and Vc could enhance the degradation effectiveness of some MCOs. Our results revealed that ceruloplasmin (CP), which carries more than 95% of the total copper in healthy human plasma ^23-27^, could mediate the degradation of Aβ and the presence of Vc could enhance the Aβ cleavage. Moreover, we found that ascorbate oxidase (AO) ^21, 28^, a widely existing MCO in healthy vegetable diet, could effectively degrade Aβs. In addition, our induced pluripotent stem (iPS) cell studies and electrophysiological analysis with brain slides suggested that AO could attenuate Aβ induced neurotoxicity and deficit in synaptic transmission in the hippocampus. To the best our knowledge, it is demonstrated for the first time that MCOs can induce degradation of peptides/proteins.

## Results

### Discovery of the degradation of FAM-Aβ42 by CP and Vc

Aβ peptides have been intensively investigated for seeking therapeutics and diagnosis of AD. Mounting evidence suggests that timely clearing of Aβ in the brain plays a critical role in AD prevention, because the clearance can prevent the accumulation of high concentrations of Aβ species, and consequentially avoid the vicious effects of the amyloidosis on brain functions ^29-34^. It is well-known that an equilibrium of Aβ exists between the brain and peripheral systems, and the timely clearance of Aβ in the blood can prevent the over-accumulation of Aβs in brains ^35-37^. In our previous report, we demonstrated that Aβs could be degraded by copper (II) in the presence of Vc ^22^. To investigate whether the copper/Vc-mediated degradation can be observed in blood, we first performed control experiments with plasma without externally added copper salts. Surprisingly, we found that FAM-Aβ42, a fluorescent dye conjugated Aβ, could be degraded by the plasma (Figure 1a,b). This unexpected result prompted us to speculate that the cleavage could be related to copper-containing proteins/enzymes. In this regard, ceruloplasmin (CP) caught our attention, due to the fact that it carries more than 95% of the total copper in healthy human plasma ^24^. To validate our speculation, we incubated FAM-Aβ42 with CP or CP/Vc for 48 hours, and the mixtures were then subjected to SDS-PAGE gel electrophoresis. Indeed, we observed one degraded band (band A in Figure 1c) that migrated faster than the monomeric bands on the gel in CP and CP/Vc groups. This result was similar to the positive control experiment with Cu(II)/Vc (Figure 1c). Quantification of the total ROI signal from bands A in Figure 1c indicated that CP could induce the cleavage of Aβ and the presence of Vc could enhance the level of cleavage fragment (25% increasing) (Figure 1d).

**Figure 1.**
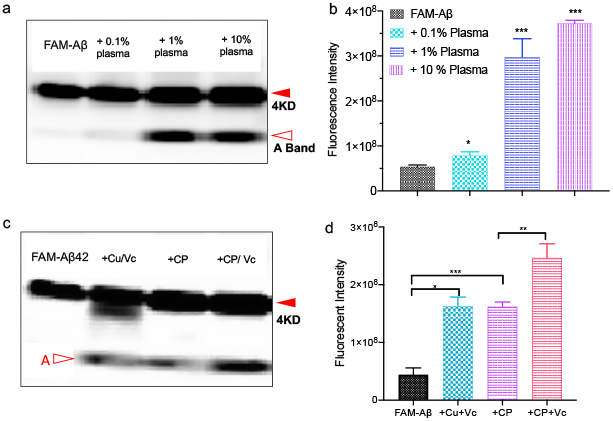
Discovery of Ceruloplasmin-induced Aβ degradation. (a) SDS-PAGE of FAM-Aβ42 with different concentrations of plasma. Fast migrating bands (A band) can be clearly seen. (b) Quantification of band A in (a) (n = 6). (c) SDS-PAGE of FAM-Aβ42 in the presence of Cu/Vc (positive control), Ceruloplasmin (CP), CP/Vc. The degradation product of FAM-Aβ42 from CP and CP/Vc groups showed similar mobility as the positive control group. (d) Quantification of band A in (c) (n = 4). P values: * P < 0.05, ** P < 0.01, and *** P < 0.001

In our previous study, we identified that FAM-Aβ1-6 was the major cleaved product in the presence of Cu(II) and Vc, and this fragment had faster mobility than FAM-Aβ42 monomers (Figure 1c). Similarly, the cleaved band from CP/Vc group showed the same mobility as the Cu(II)/Vc group, suggesting the major degradation product from the CP/Vc group is FAM-Aβ1-6.

To investigate the efficiency of the cleavage with CP and the combination of CP and Vc, we incubated FAM-Aβ42 with different ratios of CP varying from 1:0.1 to 1:5. For quantification, we used the total region-of-interest (ROI) signal from bands A (Figure S1a) and normalized the values to total ROI signals from the same positions of bands A in the control FAM-Aβ42 lane. The results indicated that the ratio was critical for efficient degradation, with a high CP ratio promoting more efficient cleavage. The presence of Vc in each group indeed has an enhanced effect, and the concentration-dependent trend in the presence of Vc was consistent with the results in the CP-alone groups (Figure S1b). However, albumin, which is the main protein of human blood plasma but does not contain copper, could not induce this degradation of FAM-Aβ42 (Figure S1c), suggesting copper inside CP plays an important role in the Aβ degradation.

To investigate whether other well-known anti-oxidants can promote the degradation of FAM-Aβ42, we tested exogenous vitamin E (Ve), curcumin, resveratrol, endogenous dopamine, and norepinephrine as reducing reagents. Different from our previous study with Cu^2+^/Vc, we found that neither of these anti-oxidants could enhance the degradation of FAM-Aβ42 compared with CP only group (Figure S2a).

### Validation of CP/Aβ interaction via fluorescence spectral studies, fluorescence anisotropy, and single-molecule TIRF imaging

To validate the interaction between CP and Aβ, we first monitor the fluorescence intensity changes of FAM-Aβ42 during incubation. Because Aβs have a high tendency to aggregate in PBS solutions, the fluorescence intensity of FAM-Aβ42 itself is gradually decreasing due to the aggregation-induced fluorescence quenching (Figure 2a). However, in the presence of CP, the decrease was considerably attenuated (Figure 2b). To investigate the strength of the interaction between Aβ42 and CP, fluorescence polarization (anisotropy) assay (FPA), a homogeneous method that provides rapid and quantitative analysis of diverse molecular interactions, was used. In this regard, we monitored the fluorescence polarization (FP) values of FAM-Aβ42 in the presence of increasing concentrations of CP. We found that the apparent Kd for the Aβ42/CP interaction was 122.7 ± 6.7 nM (Figure 2c), suggesting that the binding between Aβ42 and CP is strong.

**Figure 2.**
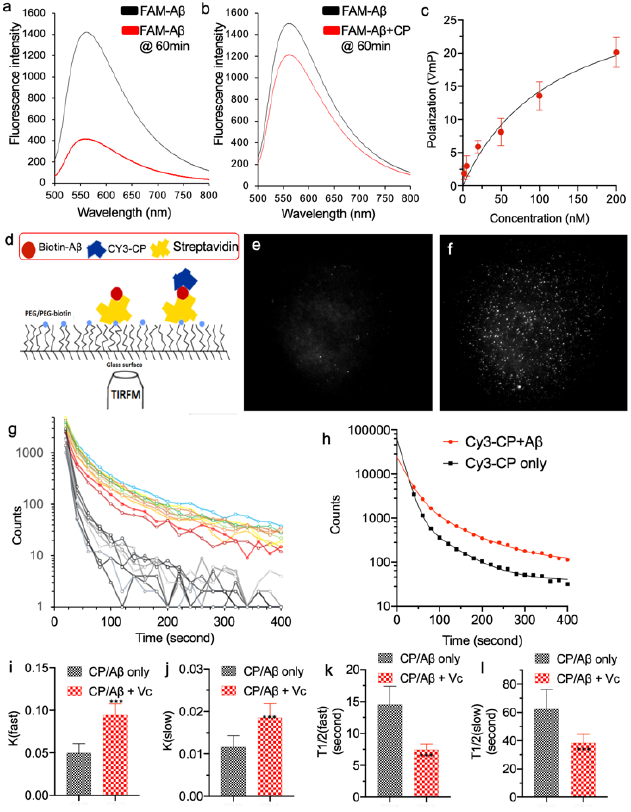
Validation of the interaction between CP and Aβ. (a) Fluorescence spectra of FAM-Aβ42 at 0 minutes (black line) and 60 minutes after incubation in PBS buffer. An apparent intensity decrease can be observed, which is due to the Aβ42 aggregation-induced fluorescence quenching. (b) Fluorescence spectra of FAM-Aβ42 at 0 minutes (black line) and 60 minutes in the presence of CP after incubation in PBS buffer. The intensity decrease was considerably attenuated due to the presence of CP. (c) Fluorescence polarization anisotropy (FPA) assay of FAM-Aβ42 with different concentrations of CP. (d) Diagram of TIRF imaging with Aβ42, which was biotinylated to be immobilized on a slide, and Cy3-conjugated CP. (e-f) TIRF images of Cy3-CP only (e), and Cy3-CP/Aβ42 (f). Apparent stronger signals could be observed from the CP/Aβ42 group. (g) Time-course of TIRF imaging of 10 trials of each group.

To further confirm the binding between Aβ42 and CP at the single-molecule level, we used a total internal reflection fluorescence microscope (TIRFM), under which the surface-bound fluorophores are selectively excited to emit fluorescence, while the non-bound fluorophores are not excited (Figure 2d) ^**38**^. TIRFM, an important tool of biophysics and quantitative biology, has been widely used to observe the fluorescence behaviors at the single-molecule level ^**39-41**^. To characterize the interaction, we first immobilized Aβ42 on the surface of microfluidics via biotin-Aβ42 and streptavidin, and the Cy3-labeled CP was added to each micro-channel (Figure 2d). Imaging was conducted and recorded for each channel for 10 minutes (Figure 2d). As expected, the image brightness from CP/Aβ was much higher than that from CP alone (Figure 2e,f). Fluorescence intensity quantification of multiple trials further confirmed the binding between CP and Aβ (Figure 2g). Non-linear regression fitting of the intensity indicated that CP/Aβ interaction had transient and stable binding modes (Figure 2h). The average transient K(fast) was about 0.048 and stable K(slow) was about 0.011; while the half-lifetimes of transient and stable binding were about 14.6 s and 62.3 seconds, respectively. Interestingly, we found that in the presence of Vc, both K(fast) and K(slow) significantly increased (0.092 count/s and 0.018 count/s) (Figure 2i, j). Consistently, the half-lifetimes with Vc considerably decreased (7.5 s and 39.0 s) (Figure 2k, l). The differences between with Vc and without Vc may provide explanations for the degradation enhancement in the presence of Vc. Taken together, the above results from fluorescence spectral testing, PFA assay and TIRF imaging suggested strong interactions between CP and Aβs.

### Preliminary mechanism studies for the degradation of Aβ peptides

In our previous study, we demonstrated that Cu(II) could induce Aβ degradation in the presence of exogenous anti-oxidants such as Vc and endogenous anti-oxidants such as dopamine. Since CP is the major copper-carrying protein in the blood, we hypothesized that Cu(II) in CP might play an important role in the CP-mediated degradation of Aβs. To prove our hypothesis, we incubated FAM-Aβ42 with CP and Vc in the absence or presence of EDTA, which is a strong Cu(II) chelator. Cu/Vc/EDTA was used in the control experiment. As shown in Figure S3a, the cleaved fragment of Aβ induced by copper and Vc disappeared in the presence of EDTA, indicating that the chelation of copper by EDTA inhibited the degradation. Surprisingly, we found that the presence of EDTA induced more degradation when incubated with CP or CP/Vc (Figure 3a, b). We reasoned that the chelation competition for copper between CP and EDTA could increase the availability of Aβ in a transient binding mode that is responsible for the cleavage of Aβ. To validate this speculation, we performed TIRF imaging in the presence of EDTA. Indeed, we found that the K(fast) increased, and the half-lifetimes became shorter in the presence of EDTA (Figure 3c, d). In our previous study, we reported that Diethyldithiocarbamate (DDC), an active metabolite of the anti-alcoholism drug disulfiram, could effectively coordinate with copper ^42^. In this report, we found that, similar to EDTA, DDC also could enhance the degradation (Figure 3a-d). Likewise, the K(fast) increased, and the half-lifetime of the transient mode decreased in the case of with Vc (Figure 3d). Taken together, all of the data indicated that the copper in CP played an important role in the Aβ degradation.

**Figure 3.**
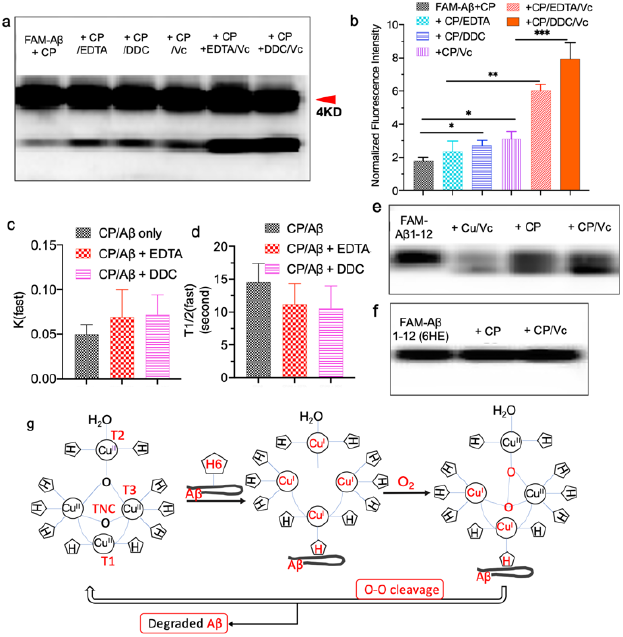
Preliminary mechanism studies of CP-induced degradation of Aβs. (a) SDS-PAGE of FAM-Aβ42 with CP and in the presence of EDTA, DDC, and Vc. (b) Quantification of the A bands in (a). (c-d) Quantification of K(fast) and T1/2(fast) from TIRF images in the presence of EDTA and DDC. (e) SDS-PAGE gel of FAM-Aβ1-12 in the presence of Cu/Vc, CP, and CP/Vc. (f) SDS-PAGE gel of FAM-Aβ1-12 with a mutated sequence (6HE) in the presence of Cu/Vc, CP, and CP/Vc. No cleaved band could be observed, suggesting H6 is crucial for the CP-induced degradation. (g) A proposed speculative mechanism for the MCO-induced degradation of Aβ peptides (H = histidine). Aβ binds to T1 copper in the oxidized form via Cu-coordination of H6 of Aβ. P values: * P < 0.05, ** P < 0.01, and *** P < 0.001.

Our previous study and other groups reported that Cu(II) could coordinate with three imidazoles in the Aβ peptide (H6, H13 and H14) ^43-49^, and we found that amide bond near the histidine could be cleaved in the presence of copper and antioxidants ^22^. In this report, to investigate whether the CP-copper has the similar binding modes as free copper, we incubated CP with several Aβ peptides and its fragments. First, we conducted tests with scrambled FAM-Aβ42, no cleaved band was observed after incubating it with CP and CP/Vc (Figure S3b), indicating that the arrangement of histidines has significant influence on the interaction between CP and Aβ.

To further elucidate the degradation site, we performed degradation experiments with Aβ fragments. Apparent cleavage bands with FAM-Aβ1-12 could be observed (Figure 3e), while no cleavage band with FAM-Aβ11-42 (Figure S3b), indicating the H6 is necessary for the interaction between CP and Aβ. This is also consistent with our previous report. Moreover, no cleavage was observed in mutated FAM-Aβ1-12 after replacing the H6 with Glutamic acid (H6E) (Figure 3f), which further confirmed the essential role of H6. However, although both FAM Aβ6-14 and FAM-Aβ10-14 have histidines in the fragments, no cleavage band with either FAM-Aβ6-14 or FAM-Aβ10-14 was observed (Figure S3c), suggesting that Aβ1-12 is essential for the binding to CP.

Based on the above results, we proposed a speculative mechanism for MCO-induced degradation of Aβ peptides. In MCOs, copper ions belong to three types, i.e, Type 1 (T1) accepts substrates and electrons, while Type 2 (T2) and Type 3 (T3) form the trinuclear center (TNC) that is the active site for O_2_ reduction ^16, 19^-^21, 25^-^27, 50^. In this study, Aβ binds to T1 copper of the oxidized form of an MCO via H6 (histidine 6) coordination and consequentially reduces Cu(II) to Cu(I) form, which is ready to accept O_2_ to form the peroxy intermediate. During the cleavage of the O-O bond of the peroxy intermediate, the bound Aβ peptide is degraded into pieces, and the MCO returns back to its oxidized form (Figure 3g). Our TIRF imaging data indicated that CP/Aβ binding had a transient mode and a stable mode, and Vc, EDTA, DDC could altered the kinetics of signals. The decay of signals could be attributed to off-binding and the degradation of Aβs. It is likely that Vc participated electron transfer from T1 to TNC, while ETDA and DDC lowered the redox potential of TNC to accelerate the turnover rate of CP. However, intensive investigation is needed to elucidate the mechanism in detail.

### Degradation by ascorbate oxidase and other multiple-copper oxidases

Considering the relevance of copper and the redox nature of the degradation, we speculated that the degradation could be a more general feature for copper-containing proteins, particularly for multi-copper oxidases (MCOs). Bilirubin oxidase, an endogenous protein, is a copper-containing polyphenol oxidase that shares similar copper coordination structure CuSS*N2 (S = Cys, S* = Met, N = His) as the ceruloplasmin family ^13, 50-52^. As expected, we observed faint cleaved bands from the mixture of FAM-Aβ42 and bilirubin oxidase (BO) (Figure 4a). To investigate whether exogenous MCOs have similar effects on the degradation, we paid attention to Ascorbate oxidase (AO) ^21^, a widely existing enzyme from a variety of vegetables. Remarkably, the effect of AO on degradation was much stronger than that of CP or BO. We observed more than 50% cleavage of FAM-Aβ42 after 4 hours of incubation (Figure 4b,c). Interestingly, no significant degradation enhancement was observed when AO was incubated with Vc (Figure 4a,b, and Figure S4a). This is likely due to the competition between Vc and Aβ for AO consumption. Since AO is widely available from vegetables, it is important to investigate whether high temperature can inactive its degradation function. As expected, AO lost its degradation capacity if it was heated at 75ºC for 2 hours (Figure S4b), which is consistent with other reports ^53^. To investigate whether AO-mediated cleavage is O_2_ dependent, we bubbled N_2_ gas into the incubation buffer. Indeed, the cleavage was considerably reduced when the samples were bubbled N_2_ gas (Figure 4e,f). Interestingly, we found that another copper-containing protein, Keyhole limpet hemocyanin (KLH) ^54^, also could induce the Aβ degradation in the presence of Vc (Figure S4c).

**Figure 4.**
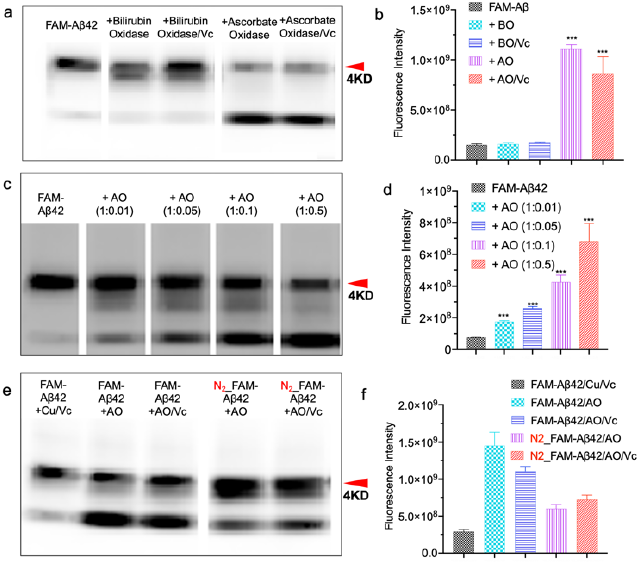
Degradation of FAM-Aβ42 with other copper-containing proteins. (a) SDS-PAGE of FAM-Aβ42 with Bilirubin Oxidase (BO), BO/Vc, Ascorbate Oxidate (AO), AO/Vc. (b)Quantification of images in (a). (c) SDS-PAGE of FAM-Aβ42 with different concentrations of AO. (d) Quantification of images of (c). (e) SDS-PAGE gel of FAM-Aβ42 with AO and AO/Vc with the purging of N2 gas before incubation. (f) Quantification of images in (e). P values: * P < 0.05, ** P < 0.01, and *** P < 0.001.

### AO attenuates the neurotoxicity of Aβ oligomers in SH-SY5Y and iPS neuronal cells

Since our in vitro solution tests revealed that AO was the strongest MCO for degrading Aβs, we used AO to investigate whether it can alleviate Aβ-induced neurotoxicity in cells. First, we incubated AO, Aβ42 oligomers with SH-SY5Y neuronal cells, and apoptosis activity was measured with ApopTag assay. Compared to the control group, the level of cell apoptosis was significantly reduced in the AO group (Figure 5a-d). Since SH-SY5Y cell line is derived from neuroblastoma, it does not represent the normal neuronal cells in brains. In this regard, we used mature induced pluripotent stem (iPS)-induced neuron cells, which were differentiated from blood stem cells. Tuj-1, a neuronal class III β-tubulin marker ^55^, was used to characterize the neuronal cells. After seeding for 12 days, abundant neuritic continuity could be observed from the neurons. After the mature neurons were treated with Aβ oligomers for 24 hours, apparent broken neuritic segments could be easily identified. As expected, no apparent broken segments could be observed when AO was mixed with Aβ oligomers (Figure 5e,f). Taken together, our data suggested that AO could rescue cell apoptosis and neuron toxicity that was induced by Aβ oligomers.

**Figure 5.**
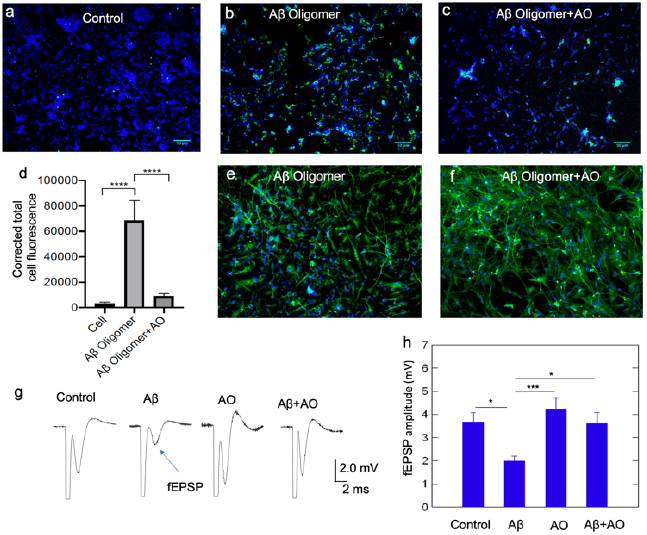
Neurotoxicity attenuation effects of AO. (a-c) cell apoptosis of SH-SY5Y cells treated with Aβ oligomer and Aβ oligomer /AO. (a) Cell only without treatment. (b) SH-SY5Y cells were treated with Aβ Oligomer (20 μM); (c) SH-SY5Y cells were treated with Aβ oligomer (20 μM) and AO (2 μM). Blue: DAPI staining (Green): immunoperoxidase staining which indicates the level of cell apoptosis. (d) Quantitative analysis of apoptosis signals for the control group (cell only), the Aβ oligomers group and Aβ oligomer/AO group. (e-f) Immunocytochemical analysis of Applied StemCell Neurons. (e) Neurons were treated with Aβ Oligomer (20 μM); (f) Neurons were treated with Aβ Oligomer (20 μM) and AO (2 μM). Green: neuronal marker Tuj-1 (neuronal type β III tubulin); Blue: DAPI. Note: Aβ Oligomer treatment resulted in a loss of neuritic continuity in neurons while Aβ Oligomer/AO treatment group showed normal length neurites. (g) Ascorbate oxidase (AO) treatment prevents Aβ-induced synaptic transmission in the hippocampus. Sample recordings of fEPSPs from control, Aβ (1 μM), AO (0.1 μM) and Aβ (1 μM)+ AO (0.1 μM) treated slices. Slices were stimulated with the pulse electric currents (0.1 ms in duration) on chaffer-collateral pathway and were recorded in the CA1 stratum radiatum. (h) Statistical summary of the maximal fEPSP amplitude in the four groups. P values: * P < 0.05, ** P < 0.01, and *** P < 0.001.

### AO prevented Aβ-induced deficit in synaptic transmission

Synaptic transmission is one fundamental function of neurons, and dementia severity correlates strongly with decreased synapse density in the hippocampus and cortex ^56^. To investigate whether AO can rescue the detrimental effects induced by Aβ oligomers in synaptic transmission, we recorded field excitatory postsynaptic potential (fEPSP) in the hippocampal CA1 area from mouse brain slides. We stimulated the Schaffer collateral pathway once every 15 seconds and recorded extracellularly from the stratum radiatum. The fEPSP, which represents the summarized synaptic events from a large population of terminals, was measured as the negative deflection of the voltage trace from the baseline. We quantified the maximal fEPSPs in the control group (no Aβ), Aβ, Aβ+AO, and AO groups. Repeated measures ANOVA of the four groups showed significant differences among these groups (p < 0.005). Turkey HSD test in different group pairs showed that the maximal fEPSP amplitudes were significantly smaller in Aβ-treated slices than in control groups (control: 3.68 ± 0.39 mV; Aβ: 1.98 ± 0.21 mV, p < 0.05) (Figure 5g). Treatment with AO prevented Aβ-induced impairment in synaptic transmission. The maximal fEPSP amplitude in the Aβ-AO group (3.61± 0.49 mV) is significantly larger than in the Aβ group (1.21 ± 0.18 mV) (p < 0.01) (Figure 5h). There is no statistical difference between the control group and the Aβ-AO group, suggesting a complete recovery of Aβ-induced detrimental effects to evoked synaptic transmission by AO administration (Figure 5h). AO alone did not cause significant changes to the fEPSP magnitude. Taken together, our electrophysiological analysis with brain slices suggested that AO could prevent an Aβ-induced deficit in synaptic transmission in the hippocampus.

## Discussion

Several enzymes, including metalloenzymes (IDE, NEP, MMP2, etc.) and Cathepsins, can induce degradation of Aβs ^4-6^. However, to the best of our knowledge, the degradation function of MCOs for Aβ peptides has not been explored. MCOs utilize copper ions as co-factors to oxidize a broad range of substrates and their degradation function for lignin families have been intensively investigated. Metal-coordination induced oxidative degradation of peptides/proteins have been reported ^5, 57, 58^. Theoretically, MCOs have potential to induce cleavage of peptides/proteins if they bind. However, such experimental evidence is rare. In this report, we demonstrated that, for the first time, highly abundant plasma CP could mediate the degradation of Aβs. Moreover, we also validated that other multi-copper oxidase could effectively cleave Aβs. Particularly, AO, a widely available protein from daily vegetable consumption ^53^, showed the strongest Aβ degradation capacity. In addition, our data showed that Vc could significantly enhance the MCO-mediate degradation of Aβs (except AO).

Consistent with our previous report, the primary cleavage site of the CP-mediate degradation is H6 of Aβ peptides. Single-molecule TIRF imaging data suggested that the binding between CP and Aβ existed in transient and stable modes, and Vc could shorten the lifetime of the transient binding state. This indicates that the cleavage occurs under the transient mode. Interestingly, EDTA and DDC could also shorten the lifetime of the transient state, suggesting the competition from EDTA and DDC may lead to more transient binding, which consequentially leads to more effective degradation. Since DDC is an active metabolite of disulfiram, an FDA-approved drug for alcoholism ^42, 59^, it may have the potential to utilize disulfiram to assist peripheral Aβ clearance. Nonetheless, more detailed studies are needed to clarify the degradation mechanism.

Among the tested MCOs, AO showed the strongest capability of degrading Aβs. Based on this data, we investigated whether AO could alleviate Aβ-induce apoptosis and fragmentation of neurites. Indeed, our data showed that AO could significantly reduce the apoptosis levels in SH-SY5Y cells, and prevent fragmentation of neurites in the mature iPS neurons. Synaptic transmission impairment has been observed in several AD animal models ^60^ and profound reduction in the size of synaptic responses in CA1 and frequent loss of field potentials were also found in aged transgenic mice ^61^. The impaired field potential in these AD mice could be due to the decreases in the density of presynaptic terminals and abnormal activation of ion channels on the presynaptic cells ^62^, Aβ-induced abnormal handling of calcium influx ^63, 64^ and oxidative stress ^65, 66^. We previously showed that incubation of brain slices in Aβ could impair evoked synaptic transmission in the hippocampus, as revealed by the decreased fEPSP magnitude ^22^. Here, our data confirmed that AO treatment could rescue the impaired synaptic strength that was induced by Aβ oligomers. Our data also shows that AO alone does not alter basic synaptic transmission in the hippocampus. Therefore, the rescue effects of the AO are likely through its degrading effects on Aβ. Considering that AO is widely available from vegetables and it has a good safety profile, we believe that utilizing AO for AD treatment is a potential approach.

AD is associated with multiple modifiable vascular and lifestyle-related risk factors. The results from the large preventive trial FINGER (Finnish Geriatric Intervention Study to Prevent Cognitive Impairment and Disability) study strongly suggested that dietary guidance, physical activity, cognitive training, and social activity could result in significant improvement of overall cognitive performance ^67^. In this report, CP, AO, and Vc are related to physical activity and healthy dietaries. For example, several groups reported that physical exercise could increase the level of CP in blood. AO and Vc widely exist in a variety of vegetables. Our data may provide certain explanations at the molecular level for the results from the FINGER study. Interestingly, most studies of diet have been focused on the devasting effects of high-fat diet and other factors that reduce the clearance of Aβs, our studies provide solutions to enhance the clearance and may guide general population to adapt to healthy lifestyle.

## Conclusions

In summary, in this report, we demonstrated the new functions of MCOs as Aβ degraders. Our findings not only expand the scope of copper-mediated degradation of Aβ, but also provide unique clues for AD therapeutics.

## Author Contributions

C. R. and J. Y. designed the project. J. Y., L.C., and C.C. executed gel electrophoresis experiments. J. Y. and K. R. performed TIRF imaging. C. R. and J. Y. analyzed the results and prepared the manuscript. W.M. and C. Z. offered constrcutive suggestions on this work. H.Y. performed fEPSP test. C. R. and Y. L. supervised the work. All authors contributed to the discussion and editing of the manuscript.

## Conflicts of interest

There are no conflicts to declare.

## Acknowledgments

This work was supported by NIH Grants R21AG059134 (C.R.), R01AG055413 (C.R.), and S10OD028609 (C.R.). The author J.Y. is grateful for the scholarship support from China Scholarship Council (CSC) and the grant BK20210416.

## Notes

### Competing Interest Statement

The authors have declared no competing interest.

