## Supplemental Information for "Degradation of amyloid beta species by multi-copper oxidases"

**Contents**

**Materials and Methods**------------------------------------------------------------------------Page 2-6

Gel electrophoresis for analyzing the degradation of Aβs ----------------------------------Page 2-4

TIRF imaging study --------------------------------------------------------------------------------Page 4

Cell apoptosis study -------------------------------------------------------------------------------Page 4-5

Toxicity study with iPS neuronal cells ----------------------------------------------------------Page 5

Electrophysiology and functional neural toxicity study--------------------------------------Page 5-6

Reference --------------------------------------------------------------------------------------------Page 6

**Supplemental Figures** ------------------------------------------------------------------------Page 7-15

SI Fig.1----------------------------------------------------------------------------------------------Page 7

SI Fig.2----------------------------------------------------------------------------------------------Page 7

SI Fig.3----------------------------------------------------------------------------------------------Page 8

SI Fig.4----------------------------------------------------------------------------------------------Page 8

Raw images for electrophoresis ----------------------------------------------------------------Page 9-15

### Materials and Methods

All of the chemicals were purchased from commercial vendors and used without further purification. Ceruloplasmin (Cat. No. C4519), Ascorbate Oxidase (Cat. No. A0157); KLH (Cat. No. H7011), Albumin (Cat. No. A9511), Bilirubin Oxidase (Cat. No. B0390) were purchased from Sigma-Aldrich. FAM-Aβ42 was purchased from American Peptide Company (Vista, CA, 92081), and FAM-Aβ1-12, FAM-Aβ1-12 (H6E), FAM-Aβ6-14, and FAM-Aβ10-14 were purchased from Genscript. FAM-Aβ11-42 (Cat. No. AS63347) and FAM-scrambled Aβ42 (Cat. No. AS60892) were purchased from Anaspec (Fremont, CA). Anti-β-amyloid antibody 6E10 was purchased from Biolegend (Cat. No. 803001). Anti-Ceruloplasmin antibody was purchased from Abcam (ab48614), and Anti-β-Tubulin III antibody was purchased from Sigma-Aldrich (T8660). Goat Anti-Mouse IgG H&L (Alexa Fluor® 488) was purchased from Abcam (ab150113). FluoroShield mounting medium with DAPI was purchased from Abcam (ab104139). Novex™ 4-20% Tris-Glycine Mini Gels were purchased from ThermoFisher. The SHSY-5Y cell line was purchased from ATCC. Neurons starter kit (Cat. No. ASE-9321K) was purchased from Applied StemCell. ApopTag Fluorescein In Situ Apoptosis Detection Kit (Cat. No. S7110) was purchased from Millipore Sigma. All animal experiments were approved by the Institutional Animal Use and Care Committee at Massachusetts General Hospital and Loyola University Chicago. A total of 15 C57BL/6J mice (2 months of age, Charles River Laboratories, Wilmington, MA) were used in this study. Mice were kept in the Loyola University Chicago Animal Facility under continuous care by facility technicians.

#### Gel electrophoresis for analyzing the degradation of Aβs:

*Sample preparation:*

*1. Degradation of FAM-Aβ42 induced by ceruloplasmin and Vc.* A 10 μL HFIP (hexafluoroisopropanol) solution (25 μM) of FAM-Aβ42 was added to a 1.5 mL LoBind eppendorf tube. After evaporating the organic solvent under vacuum, 4 μL DMSO was added to the tube, followed by the addition of 8 μL of ceruloplasmin solution in PBS (31.25 μM) and 4 μL of Vc solution (1.25 mM) and 4 μL of PBS. The resulting mixture was incubated at 37°C for 24 hours and was then subjected to gel electrophoresis.

2. *CP concentration-dependent study.* A 10 μL HFIP (hexafluoroisopropanol) solution (25 μM) of FAM-Aβ42 was added to a 1.5 mL LoBind Eppendorf tube. After evaporating the organic solvent under vacuum, 4 μL DMSO was added to the tube, followed by the addition of 8 μL of ceruloplasmin solution in PBS (3.125, 15.6, 31.25, and 156 μM) and 8 μL of PBS. The resulting mixture was incubated at 37°C for 24 hours and was then subjected to gel electrophoresis.

3. *Vc concentration-dependent study.* A 10 μL HFIP (hexafluoroisopropanol) solution (25 μM) of FAM-Aβ42 was added to a 1.5 mL LoBind Eppendorf tube. After evaporating the organic solvent under vacuum, 4 μL DMSO was added to the tube, followed by the addition of 8 μL of ceruloplasmin solution in PBS (31.25 μM) and 4 μL of Vc solution (62.5 μM, 1.25 mM, 6.25 mM, and 12.5 mM) and 4 μL of PBS. The resulting mixture was incubated at 37°C for 24 hours and was then subjected to gel electrophoresis.

4. *Degradation of other FAM labeled Aβs induced by CP and Vc.* A 10 μL HFIP (hexafluoroisopropanol) solution (25 μM) of FAM-scramble Aβ42 or FAM Aβ11-42 was added to a 1.5 mL LoBind Eppendorf tube. After evaporating the organic solvent under vacuum, 4 μL DMSO was added to the tube, followed by the addition of 8 μL of ceruloplasmin solution in PBS (31.25 μM) and 4 μL of Vc solution (1.25 mM) and 4 μL of PBS. The resulting mixture was incubated at 37°C for 24 hours and was then subjected to gel electrophoresis. For other Aβ fragments: A 8 μL dd water solution (25 μM) of FAM Aβ1-12, FAM Aβ1-12 (H6E), FAM Aβ10-14 or FAM Aβ6-14 was added to a 1.5 mL LoBind Eppendorf tube respectively. Then 8 μL of ceruloplasmin solution in PBS (25 μM) and 4 μL of Vc solution (1 mM) were added to the tube. The resulting mixture was incubated at 37°C for 24 hours and was then subjected to gel electrophoresis.

5. *Degradation of FAM Aβ42 induced by CP and Vc in the presence of EDTA and DCC.* A 10 μL HFIP (hexafluoroisopropanol) solution (25 μM) of FAM-Aβ42 was added to a 1.5 mL LoBind Eppendorf tube. After evaporating the organic solvent under vacuum, 4 μL DMSO was added to the tube, followed by the addition of 8 μL of ceruloplasmin solution in PBS (31.25 μM), 4 μL of Vc solution (1.25 mM) and 4 μL of EDTA (625 μM). The resulting mixture was incubated at 37°C for 24 hours and was then subjected to gel electrophoresis.

6. *Degradation of FAM-Aβ42 induced by other proteins.* A 10 μL HFIP (hexafluoroisopropanol) solution (25 μM) of FAM-Aβ42 was added to a 1.5 mL LoBind Eppendorf tube. After evaporating the organic solvent under vacuum, 4 μL DMSO was added to

the tube, followed by the addition of 8 μL of bilirubin oxidase solution or ascorbate oxidase or keyhole limpet hemocyanin or albumin in PBS (31.25 μM) and 4 μL of Vc solution (1.25 mM) and 4 μL of PBS. The resulting mixture was incubated at 37°C for 24 hours and was then subjected to gel electrophoresis.

*Gel electrophoresis:* Samples were separated on 4–20% gradient Tris-glycine mini gels (Invitrogen). The images were acquired on IVIS®Spectrum (Perkin Elmer) with excitation at 465nm, and emission at 520nm.

**TIRF imaging study.** This study was performed by following the published protocol[1] Briefly, basic slide passivation was performed using 5kD polyethylene glycol (PEG)+2.5% 5kD biotin-PEG (LaysanBio, MPEG-SVA-5000; Biotin-PEG-SVA-5000) in a 'clouding-point' solution on amino-silanized slides for 4 hours as described. After passivation, slides were assembled into microfluidic reaction chambers. The passivated slide channels were incubated with 0.2 mg/ml streptavidin and cell extract for 1 minute and rinsed with TBS for 3 times. Then the injection ports of the flow cell were sealed with ~0.5 μL mineral oil to prevent loss with passivation due to drying. Then the channels were coated with 200 nM biotinylated-Aβ42 in coating buffer (2 mg/ml β-globulin in TBS buffer) and washed 1 time with TBS. Then the channels were flushed with the imaging buffer (2 mg/ml β-globulin in TBS buffer) containing 100 nM CP-Cy3 or 100 nM CP-Cy3 with 250 μM Vc or 100 nM CP-Cy3 with 125 μM EDTA or 100 nM CP-Cy3 with 125 μM DDC. The surface was imaged immediately with a Nikon Ti TIRF microscope equipped with a 561nM laser at 50mW. Time series were acquired at 20sec/frame for 10 minutes with exposure time at 100 ms.

**Cell apoptosis study**

1. *Preparation of Aβ42 oligomers.* 1.0 mg of Aβ42 peptide (HFIP) was suspended in 221 μL HFIP to make a 1mM stock solution. Then a 20 μL stock solution of Aβ42 was added to a 1.5 mL LoBind Eppendorf tube. After evaporating the organic solvent under vacuum, 4 μL DMSO was added to the tube. The tube was ultrasonicated for 5 min, followed by the addition of 200 μL of F12 media. The resulting mixture was incubated at 4°C for 24 hours. Then the resulting solution

was centrifuged at 14000g for 10min at 4°C and the supernatant was removed and stored in a new LoBind Eppendorf tube as Aβ42 oligomers.

2. *Fluorescein in situ apoptosis detection Assay.* SH-SY5Y cells were planted to a 24-well culture plate with $3x10^4$ cells per well. When the cells confluency reached about 80%, the cells were treated with Aβ42 oligomers (100 µL prepared Aβ42 oligomers and 400 µL culture media), and a mixture of Aβ42 oligomers and AO (100 µL prepared Aβ42 oligomers 100µM and 400 µL culture media containing 2 µM AO). After 24 hours, the apoptosis level of the cells was detected by using a fluorescein in situ apoptosis detection kit.

**Toxicity study with iPS neuronal cells.** Mature iPS neurons were prepared using an optimized maturation medium and supplements from Applied StemCell's by following the provided protocol in Applied StemCell's Neurons Starter Kit. When the cell population reached about 90%, the cells were treated with Aβ42 oligomers (100 µL prepared Aβ42 oligomers and 400 µL culture media) and a mixture of Aβ42 oligomers and AO (100 µL prepared Aβ42 oligomers 100µM and 400 µL culture media containing 2 µM AO). After 24 hours, the neurons were washed two times with PBS and fixed with 4% paraformaldehyde in PBS for 15 min at room temperature. After aspirating the fixative solution, the neurons were rinsed twice in PBS for 5min each and permeabilized with 0.2% triton x-100 in PBS for 10min at room temperature. After aspirating the triton x-100, the neurons were rinsed twice in PBS. Then the neurons were incubated with 1% BSA in PBST for 1 hour at room temperature. After aspirating the BSA solution, the neurons were incubated with monoclonal anti-β-Tubulin III antibody at 4°C overnight. Then the neurons were rinsed three times in PBST for 5 min each and incubated with Goat Anti-Mouse IgG H&L (Alexa Fluor® 488) for 1 hour at room temperature in the dark. Then the neurons were rinsed three times in PBST for 5 min each and covered with FluoroShield mounting medium with DAPI and sealed with nail polish. Florescence images were obtained using the Nikon Eclipse 50i microscope with blue and UV light excitation channels.

**Electrophysiology and functional neural toxicity study**

1. *Tissue Preparation.* Mice were anaesthetized with isoflurane in a sealed chamber. The animals were then decapitated, and the brain was quickly removed, hemisected, and placed into ice-cold artificial cerebrospinal fluid (ACSF) for ~3 min. The cerebellum was removed, and the

brain was bisected along the midsagittal line. The superior cortex was removed, and the dorsal cortex was cut parallel to the longitudinal axis. Cyanoacrylate glue was then used to fix the brain, ventral side up, to an aluminum block. The block was secured in a Vibratome (Series 1000, Technical Products International, St. Louis, MO) and brain slices (400 μm) were obtained and incubated in ACSF at room temperature (control group). ACSF contained (mM): 120 NaCl, 2.5 KCl, 2 $CaCl_2$, 2 $MgCl_2$, 25 $NaHCO_3$, and 10 d-glucose, pH 7.4, and was continuously bubbled with 95% $O_2$ +5% $CO_2$. For the Aβ-challenged groups, slices were incubated in 1 μM Aβ. For the ascorbate oxidase (AO) treatment group, slices were incubated with 1 μM Aβ and 0.1 μM AO. As a control, the effects of AO alone (0.1 μM) were also studied.

*2. Extracellular recordings.* After 1-hour incubation, slices were moved to a recording chamber for extracellular field potential recording at room temperature. Schaffer collateral commissural projections were stimulated with a bipolar tungsten electrode (enamel-insulated nichrome wire, 125 mm diameter) for orthodromic activation of the CA1 neurons. A recording borosilicate glass electrode, filled with 150 mM NaCl, was placed in the stratum radiatum (dendritic region) of the CA1 region to record field excitatory postsynaptic potentials (fEPSPs). Constant current pulses of 0.1 ms duration (4 s interval) were generated by a Grass S88 stimulator (Grass Instruments, Quincy, MA) and delivered through an isolation unit. Pulse polarity was selected to produce the largest potentials with a clearly distinguishable artifact. Responses were filtered, amplified, and recorded with a PClamp 10 amplifier (Axon Instruments, Foster city, CA) and acquisition of data was performed using Clampex 10 (Axon Instruments).

*3. Statistics.* Throughout the text, mean ± standard error (SE) was reported. fEPSP amplitudes were given by pClamp 9 software by measuring the maximum negative deflection from the baseline. All data were verified for normal distribution and homogeneity of variance. Statistical significance was determined with one-way repeated ANOVA followed by the post hoc Turkey HSD test, using SigmaStat software (v. 3.0, Aspire Software International, Ashburn, VA). Effects were considered statistically significant at $p < 0.05$.

### Supplemental Figures

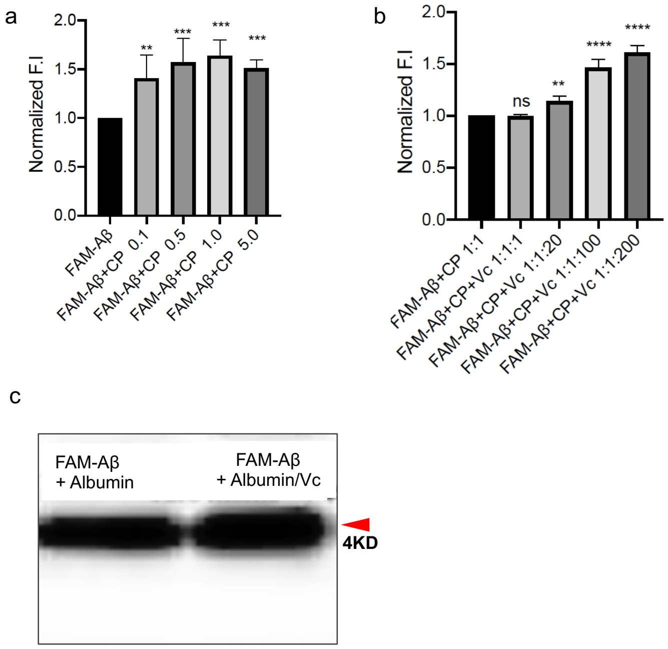


**SI Figure 1.** Degradation of FAM-Aβ induced by different concentrations of CP and Vc. (a) Quantification of SDS-PAGE of FAM-Aβ42 with different CP/Aβ ratios of 0.1, 0.5, 1.0, and 5.0. (b) Quantification of SDS-PAGE of FAM-Aβ42 with different CP/Aβ/Vc ratios of 1:1:1 to 1:1: 200. (c) SDS-PAGE of FAM-Aβ42 with Albumin and Albumin/Vc. No apparent band of degradation can be observed.

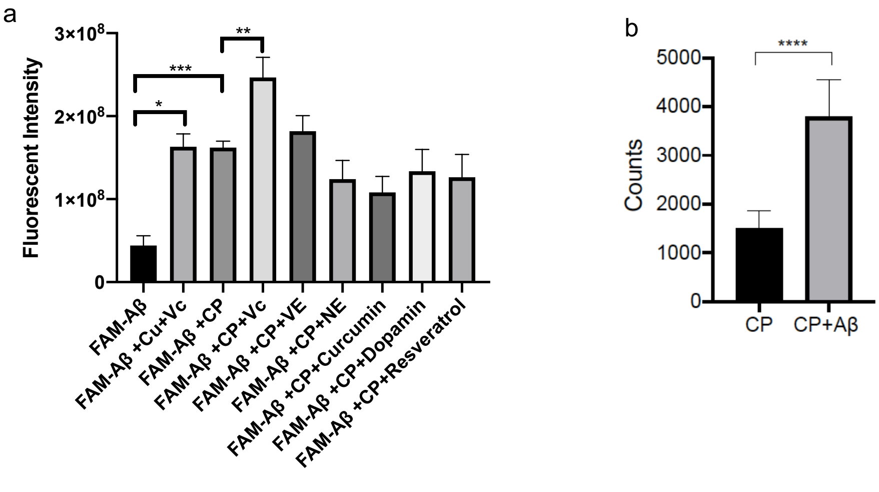


**SI Figure.2** (a) Quantification of SDS-PAGE gel of FAM-Aβ42 with CP in the presence of various anti-oxidants. b) Quantification of fluorescence intensity of TIRF images in the presence of Cy3-CP and Cy3-CP with biotinylated Aβ42.

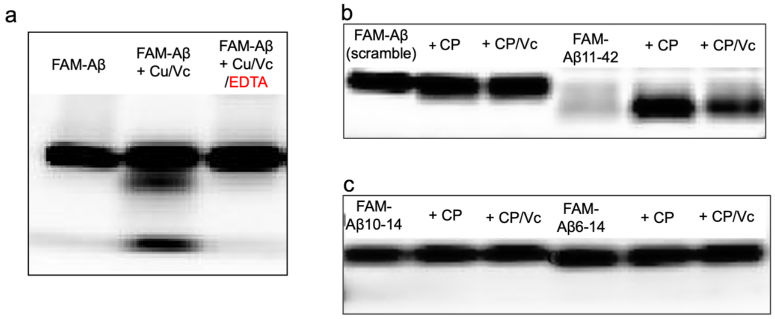


**SI Figure 3.** (a) SDS-PAGE of FAM-Aβ42 with Cu(II)/Vc (1:20), Cu(II)/EDTA/Vc (1:10:20), (b) SDS-PAGE gel of scrambled FAM-Aβ42 (Left) and fragment of Aβ (FAM-Aβ11-42) (right) in the presence of CP and CP/Vc. (c) SDS-PAGE gel of Aβ fragments FAM-Aβ10-14 (Left) and Aβ (FAM-Aβ6-14) (right) in the presence of CP and CP/Vc. No clear bands of degraded Aβs can be identified.

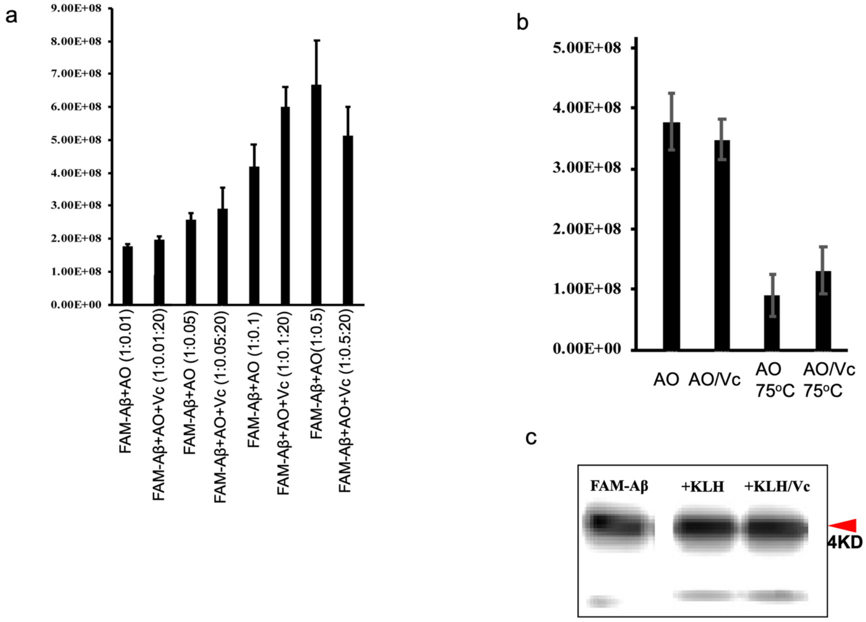


**SI Figure 4.** (a) Quantification of SDS-PAGE gel of FAM-Aβ42 with different ratios of AO in the presence of Vc. (b) Quantification of SDS-PAGE gel of FAM-Aβ42 with native AO and AO treated at 75 ºC for 2 hours. (c) SDS-PAGE gel of FAM-Aβ42 with KLH and KLH/Vc.

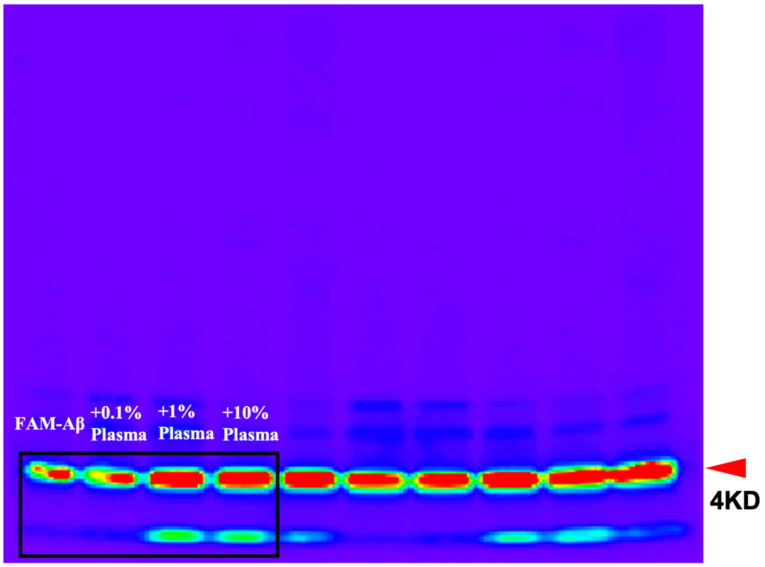


**SI Figure 5**. Raw images for Figure 1a. The data presented was framed.

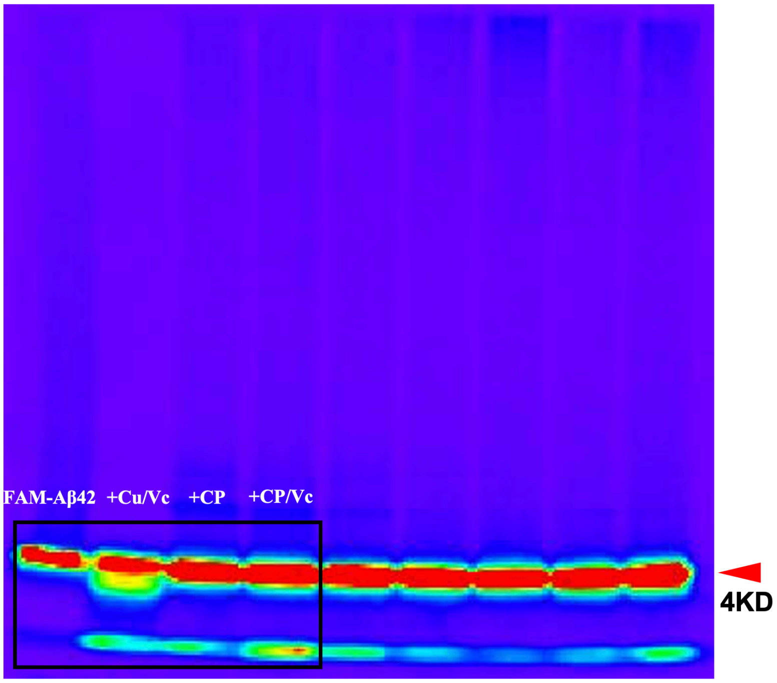


**SI Figure 6.** Raw images for Figure 1c. The data presented was framed.

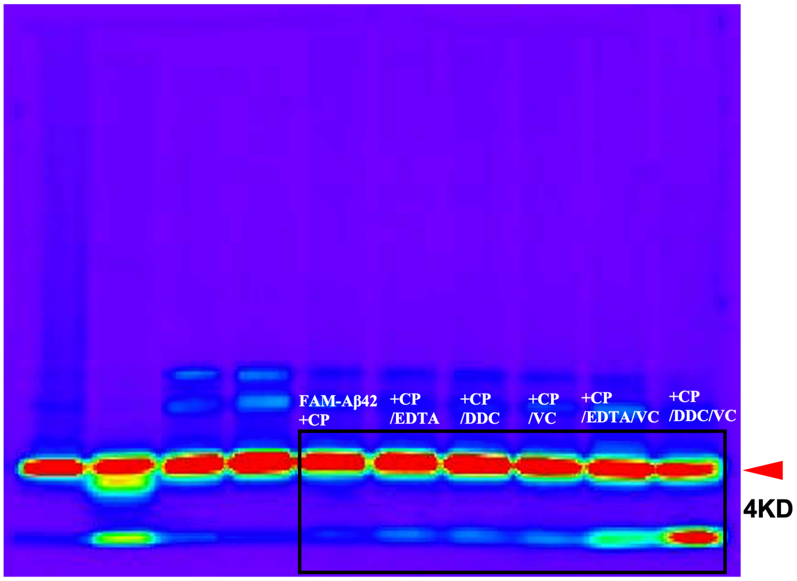


**SI Figure 7.** Raw images for Figure 3a. The data presented was framed.

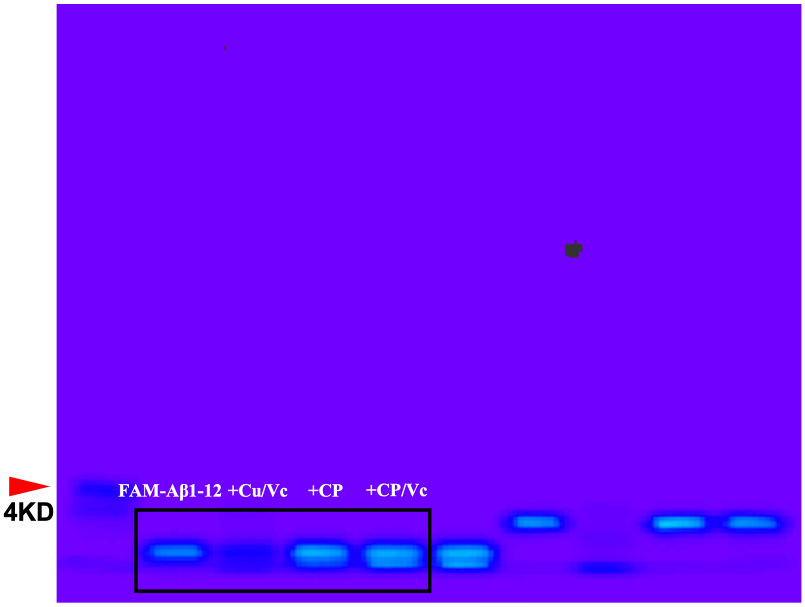


**SI Figure 8.** Raw images for Figure 3e. The data presented was framed.

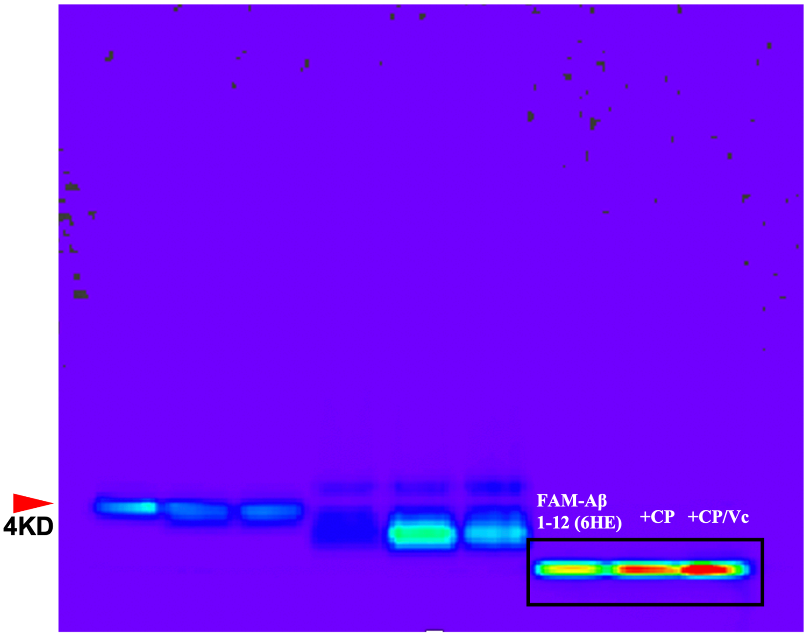


**SI Figure 9.** Raw images for Figure 3f. The data presented was framed.

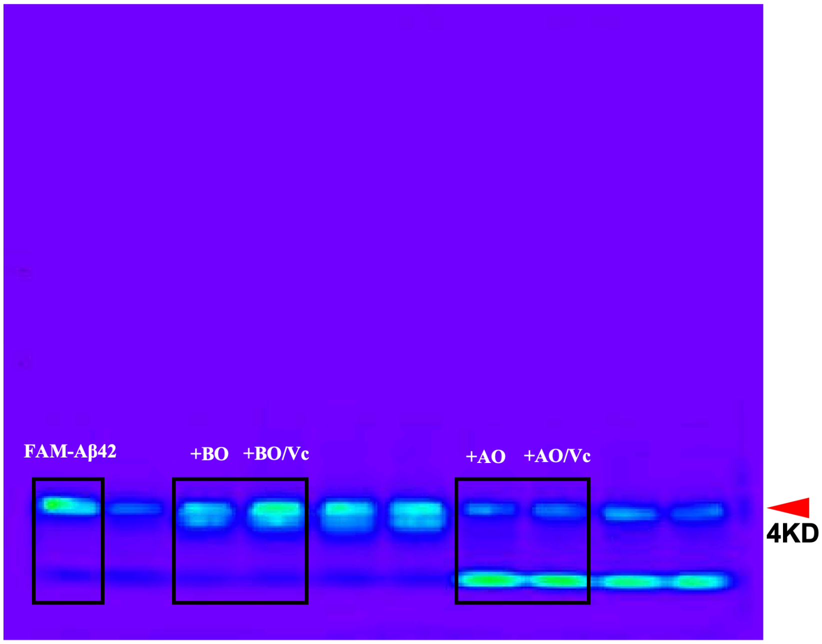


**SI Figure 10.** Raw images for Figure 4a. The data presented was framed.

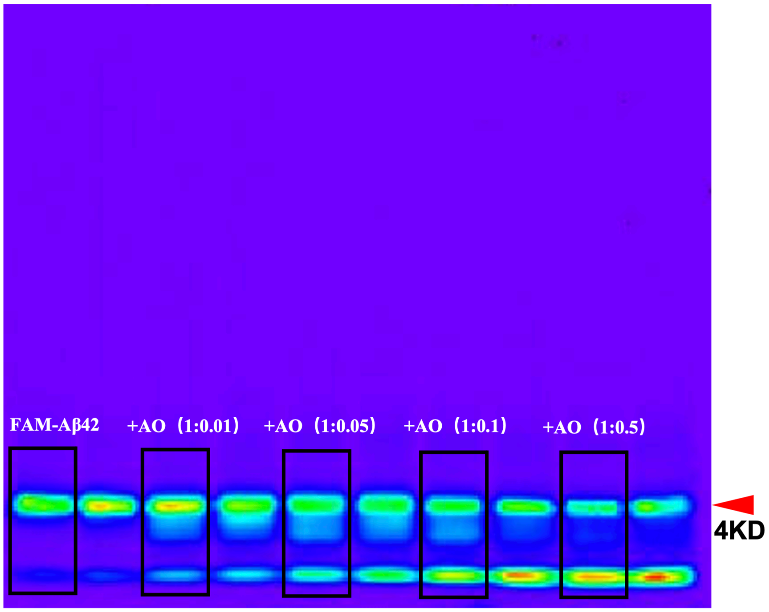


**SI Figure 11**. Raw images for Figure 4c. The data presented was framed.

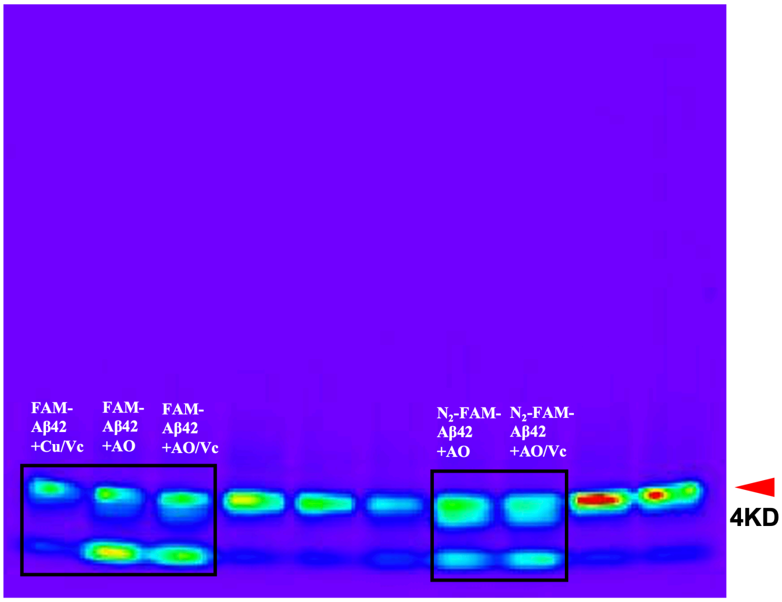


**SI Figure 12.** Raw images for Figure 4e. The data presented was framed.

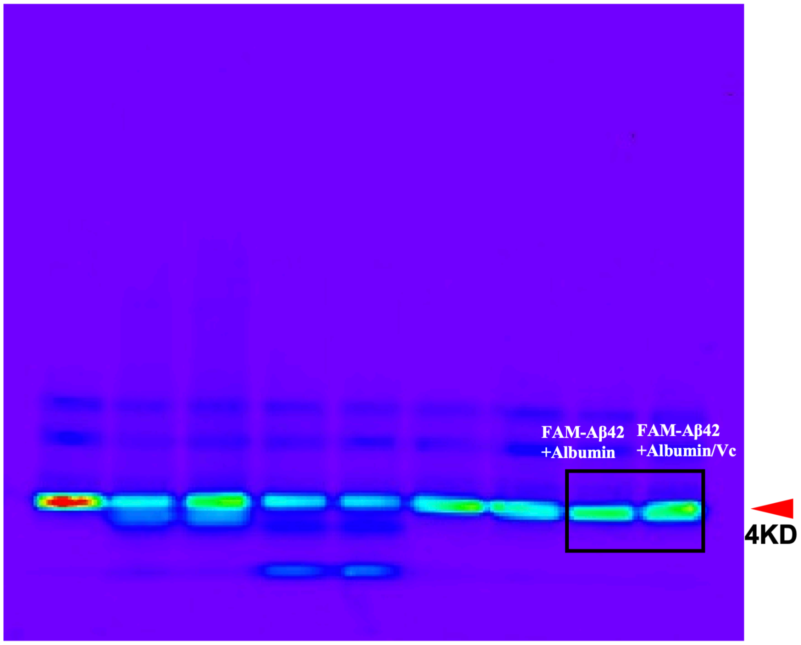


**SI Figure 13.** Raw images for Figure S1a. The data presented was framed.

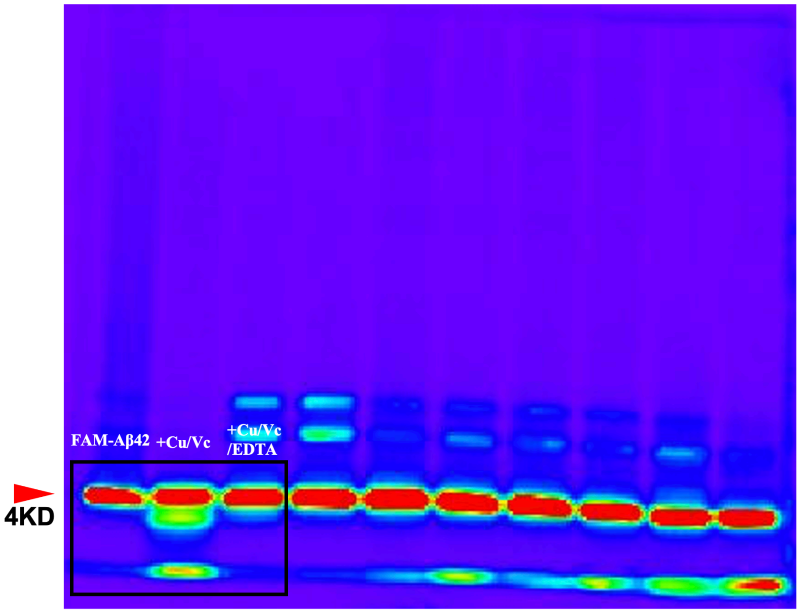


**SI Figure 14.** Raw images for Figure S3a. The data presented was framed.

`

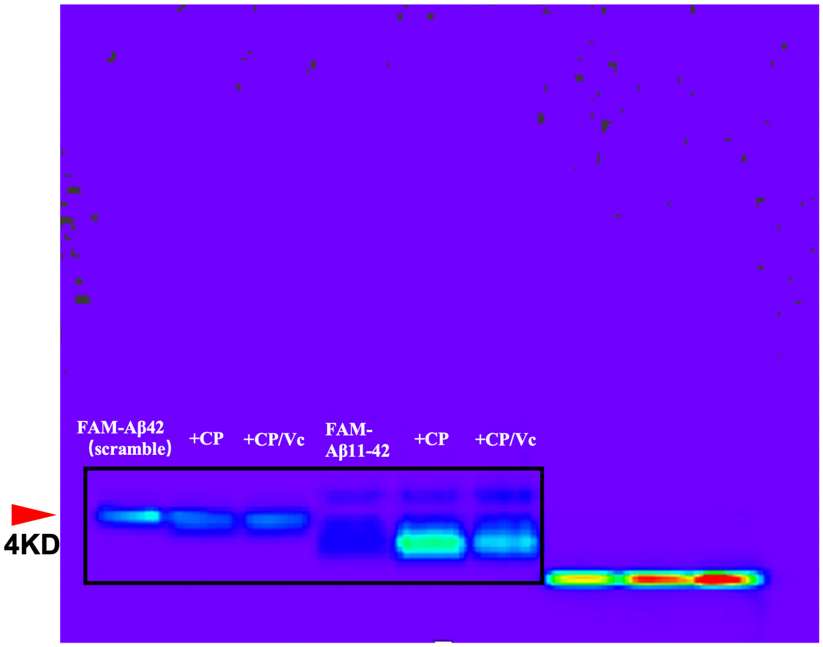


**SI Figure 15.** Raw images for Figure S3b. The data presented was framed.

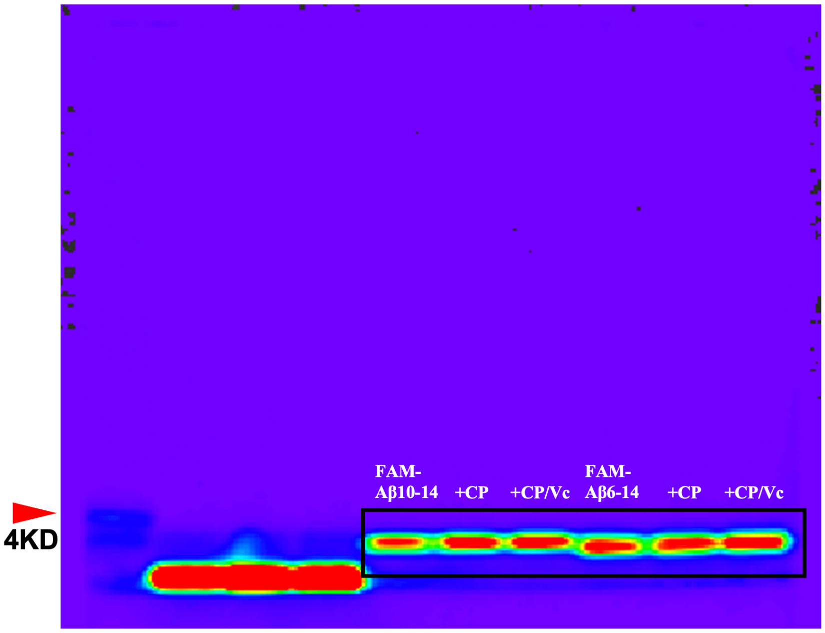


**SI Figure 16.** Raw images for Figure S3c. The data presented was framed.

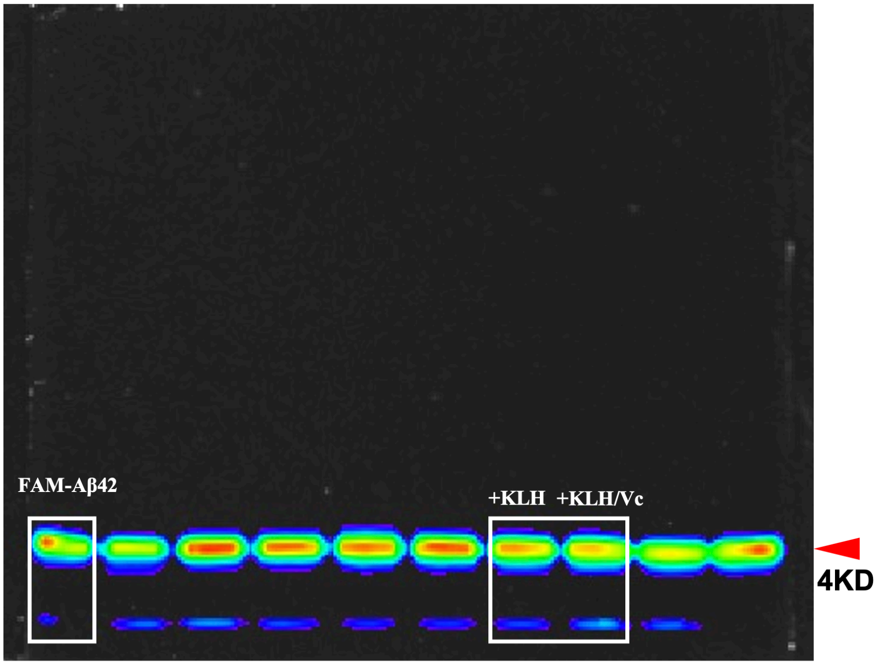


**SI Figure 17.** Raw images for Figure S4c. The data presented was framed.
